# The Sun within: active processes from two-temperature models

**DOI:** 10.1101/2023.10.21.563425

**Authors:** Faezeh Khodabandehlou, Christian Maes

**Affiliations:** Department of Physics and Astronomy, KU Leuven, Belgium

## Abstract

We propose an embedding of standard active particle models in terms of two-temperature processes. One temperature refers to an ambient thermal bath, and the other temperature effectively describes “hot spots,” *i.e*., systems with few degrees of freedom showing important population homogenization or even inversion of energy levels as a result of activation. As a result, the effective Carnot efficiency would get much higher than for our standard macroscopic thermal engines, making connection with the recent conundrum of hot mitochondria. Moreover, that setup allows to quantitatively specify the resulting nonequilibrium driving, useful in particular for bringing the notion of heat into play, and making easy contact with thermodynamic features. Finally, we observe that the shape transition in the steady low-temperature behavior of run-and-tumble particles (with the interesting emergence of edge states at high persistence) is stable and occurs for all temperature differences, including close-to-equilibrium.

## I. INTRODUCTION

Heat engines operate by extracting heat from one thermal bath at a higher temperature and delivering heat to a similar reservoir at a lower temperature, [1, 2]. Biological systems (like animal muscles, molecular motors or simple bacteria) often operate and are able to deliver useful work at a body temperature that is equal to or lower than the ambient temperature. In such a situation, when blindly applying the Carnot formula, we would predict zero, or even negative, efficiency. But obviously, those biological systems are not Carnot engines; they are active and far from the reversible limit. Nevertheless, the point of departure of the present paper is that delivering work for a two-temperature engine does not require both reservoirs to be in equilibrium, nor do we need a macroscopic or quasistatic setting. One bath can simply consist of many independent *small* nonequilibrium systems or active agents. Its corresponding temperature is then indeed only (but in some specific sense) *effective*. The involved molecules are activated and produced by chemistry, to provide the necessary fuel. Yet, the activation is not first entirely degraded into heat, after which a Carnot engine is started. That is why biological processes need to be fast and to have some persistence, exactly to avoid that thermalization.

The fact that energy transfer and the related possibility to deliver work need not be driven by differences in *real* temperature between thermal (equilibrium) baths has been noted before, *e.g*. in [3–7]; it suffices that the “effective” temperatures (*e.g*. as calculated from the relative occupations in a multilevel system) between two regions are different. The main point is that the chemical reaction brings a product (molecule or subsystem) in an excited state, with a statistical occupation of energies showing a homogenized or even inverted population with respect to the thermal distribution at the ambient temperature. That activation is often concentrated in only few degrees of freedom (a vibration mode or a couple of two-level systems), *effectively* mimicking a high-temperature spot.

As an example, the hydrolysis of an adenosine triphosphate (ATP) molecule means to split high-energy phosphoanhydride bonds with a release in chemical energy of about say 0.4 eV/molecule. Spreading that over two degrees of freedom gives an effective temperature *T*_eff_ ≃ 0.4 eV/k_*B*_ which is like 20 times room temperature (where the average thermal energy per degree of freedom equals *k*_*B*_*T/*2 = 0.01 eV); see [2] for the more detailed calculation. In other words, instead of zero or negative, a naive application of the Carnot formula would yield corresponding efficiencies reaching much higher values than we are used to for our macroscopic engines.

We thus imagine a collection of such nonequilibrium “hot spots” in contact with a heat bath at ambient (or body) temperature, creating the possibility to deliver work, [8, 9]. We have little to say here about the specific nature of those “hot spots” in their biological environment. It is not unnatural for Eukaryotes to associate the spots with mitochondria, but for bacteria, there will be other centers of activity. Surely, thermal baths are present as well, *e.g*. in the form of an internal watery medium in which the agents dissipate heat and/or outside at ambient temperature. Yet, in contrast with the usual setup with heat engines, we have various small systems (few degrees of freedom), very much out of equilibrium, coupled to a (more traditional) thermal bath such as a bloodstream, a water bag or an external environment. That setup is able to perform work, even when the small system operates in noisy surroundings where the passive elements appear to be at temperatures equal to or lower than the ambient temperature.

To continue, however, one needs a clear thermal interpretation of these models, which brings us to the second main point of the paper.

To insert the study of Life into the realm of statistical physics requires understanding it as a nonequilibrium process. It demands a quantitative description of the departure from equilibrium. Life processes typically possess a continuously varying amplitude where the biochemical fuel or driving can at least in principle be varied far enough to touch *death, i.e*., equilibrium and detailed balance. The latter may give a useful theoretical reference or point of contrast for building models mimicking life processes, *e.g*., to invoke local detailed balance leading to relevant thermal interpretations. Indeed, a problem with current models of active systems is that there is no clear reference which can be called the equilibrium point from which perturbation theory can start. Usually, one considers the passive limit (such as the asymptotics of large flipping rate for run-and-tumble particles) as the equilibrium limit. While that is formally possible, it is physically odd. After all, the flip rate is a time-symmetric parameter and does not directly represent any fueling or driving. The present paper launches two-temperature models through which important representatives of active processes get a more standard place in nonequilibrium physics.

The next Section II directly deals with models of active particles as two-temperature processes. Section II A discusses the case of run-and-tumble particles, followed by Section II B on active Brownian dynamics. For both, we propose a two-temperature interpretation. We observe transitions in the stationary behavior as a function of the system parameters such as the effective temperature, from which robustness of the so-called shape transition, [10, 11], can be derived, as our modeling allows to extend the phenomenon in the two-temperature setting. Furthermore, local detailed balance is restored and, from there, a thermal characterization becomes possible. In particular we are able to construct the (close-to-equilibrium) McLennan distribution, as recalled in Appendix A.

For modeling a “hot spot,” Section III A treats an activated molecule, in particular in the form of a molecular switch where the order of the energies in a multilevel system gets inverted at random times, and as a flashing vibrational mode in Section III B. There we show how the activation indeed raises the effective temperature, in the sense of increased uniformization of energy occupation statistics. It also provides theoretical modeling relevant in the discussion concerning hot mitochondria, [12–17].

The paper has many natural continuations, for example concerning the details of the extractability of work in living systems. See *e.g*. [18] for the method of bacterial rectification, and also muscle efficiency is discussed in many places; see *e.g*. [19]. For the biological physics of population inversion as visible from calorimetry on active systems, we refer to [20, 21]. Bringing these subjects together and unified under the governing body of nonequilibrium statistical mechanics is a theoretical contribution to which the present paper wishes to add.

## II. ACTIVE PARTICLES

Here comes the embedding of standard models of self-locomotion in a two-temperature setting, with one high effective temperature governing the dynamics of an internal variable (hot spot). Note that only in Section III we add a description of “agitated” systems to represent such “hot spots,” and we discuss the for our purpose relevant meaning of *effective* temperature.

The particles of the present section are subject to overdamped dynamics, possibly under the influence of thermal noise and a confining potential, while the “hot spot” adds another source of (colored) noise. We do not enter into the details of the mechanisms through which that effective heat engine delivers work and produces motion. The few models we discuss here can be associated to flagellar locomotion. Yet, locomotion in living material obviously does not always involve flagellar motion. There is also crawling, swimming, and gliding of bacteria, responding differently to the environment and its gradients and showing a great variety of mechanisms for employing low-entropy sources, [22]. Nevertheless, we believe the theoretical foundations of our proposal are more universal than apparent in the following (bacterial) models. In particular, the biological interpretation of the “hot spots” is ignored here, but for Eukaryotes it is certainly tempting to suppose that they are represented by “hot mitochondria”, [12–17].

### A. Run-and-tumble particles

Run-and-tumble particles (RTPs) are among the best-known toy models of active matter. They share important features with bacterial locomotion, with *E. Coli* as the main representative. We refer to the by now many studies and reviews, including [23–25].

For simplicity we consider position *x* ∈ ℝ subject to the dynamics,

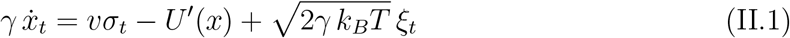

where soon we will put *γ* = 1, *k*_*B*_ = 1. The potential *U* is spatially confining around the origin. The *ξ*_*t*_ is standard white noise with ambient temperature *β*^*−*1^ = *k*_*B*_*T* giving it strength. The hot spot dynamics is represented by *σ*_*t*_ = ±1, which flips sign

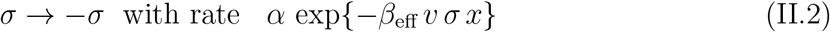

depending on the position *x* of the walker. For *β*_eff_ = 0, the *σ*_*t*_ represents standard dichotomous noise, and then (II.1) (often with *T* = 0) describes a one-dimensional run-and-tumble motion with tumbling rate *α*. Jointly, in any event, (*x*_*t*_, *σ*_*t*_) is a Markov process with one continuous and one discrete variable, respectively, coupled to the heat bath at temperature *T* and the *internal* heat bath at effective inverse temperature *β*_eff_. Fig. 1 gives an impression of the situation we wish to imagine more generally.

**FIG. 1:**
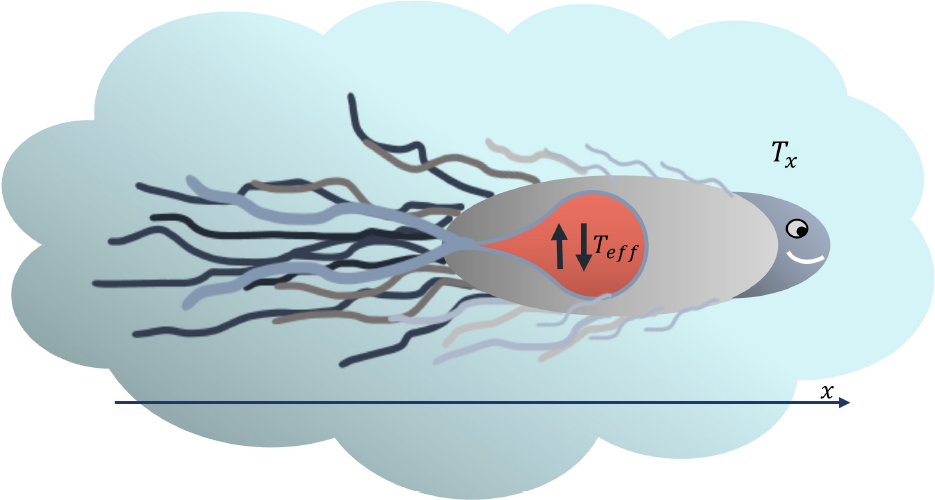
Cartoon of an active particle, for zooming in on the “hot spot” at effective temperature *T*_eff_, coupled to the locomotion system at ambient temperature *T* = *T*_*x*_, with *x* the position of the overdamped swimmer. The biological interpretation of the “hot spots” will certainly differ per organism. For *E. Coli* as modeled via RTPs in the present section, we have in mind motility proteins (the rotary engine) with power provided by a proton flow, while for Eukaryotes, it is tempting to think about mitochondria.

At first sight, there is some problem with the tumbling rates (II.2), as they depend on the location of the spatial origin. That is however the same origin as already specified by the confining potential. Moreover, from bacterial chemotaxis, spatial gradients in flipping rate are not uncommon. Furthermore, for *β*_eff_ ≃ 0 (high internal temperature) that dependence becomes very weak, and for the locomotion part (II.1), the force *v σ*_*t*_ is fully ignorant about the position. There is also something to gain.

The great advantage of the parametrization in (II.1)–(II.2) is that we have a clear reference *β*_eff_ = *β* (keeping *α* ≠ 0) where the condition of detailed balance holds. Fixing *α >* 0, we can now interpolate between the purely random flipping (*β*_eff_ = 0) and the equilibrium dynamics (*β*_eff_ = *β*) via that temperature-driven analog. At the same time, it brings the RTP–modeling within the realm of local detailed balance [26] from where thermal features are more easily interpreted. There is a contact between (effectively) two thermal baths, one at inverse (ambient) temperature *β*, and one at inverse (internal but effective) temperature *β*_eff_. The difference *β* − *β*_eff_ in inverse temperatures provides the driving, much as in a traditional heat engine. In that way, we interpret the motion of RTPs as originating from a two-temperature model, from which the usual thermodynamic interpretation of heat flux holds; see *e.g*. [27]. We work therefore under the questions of the papers [28, 29]. Other effective thermal descriptions exist of course, as *e.g*. in [30].

#### 1. Shape transition

The internal bath is given effective temperature *T*_eff_, imagining internal active processes carried by low-dimensional degrees of freedom (see Section III). Here we look at the dependence on *β*_eff_ for the stationary dynamics and static fluctuations. In particular, we are interested in observing the shape transition between an edgy and a confined regime [10, 11], as a function of the effective temperature.

The process satisfies detailed balance for *β*_eff_ = *β* (equal temperatures), with equilibrium density

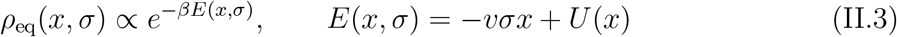

at (unique) inverse temperature *β* (independent of *α*, as it should), at least as long as the potential *U* is sufficiently confining. For example, for the harmonic potential *U*(*x*) = *x*^2^, we have

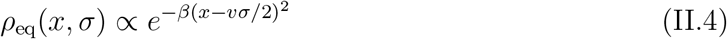

and it depends on the propulsion speed *v* and on *β* whether *ρ*_eq_(*x*) = (*ρ*_eq_(*x*, +1) + *ρ*_eq_(*x*, −1))/2 has its maximum at the origin *x* = 0 (confined regime) or at the edges *x* = ±*v/*2 (edgy regime).

We are interested mostly in large *β* (small thermal noise for locomotion).

Suppose 0 *< β*_eff_ ≤ *β*: the flipping rate *α* exp{−*β*_eff_ *v σ x*} is going to be small when *σ x >* 0 and *β*_eff_ is large, and hence, say for *σ* = 1, the particle will continue to large *x* > 0 without flipping (flipping rate decreasing ever more). For example, for *σ x* ∼ *v*, the particle is expected to get trapped for a long time, especially at large propulsion speed *v*, and *α* needs to be truly large to avoid that. For example, it remains true at *T* = 0, from (II.1), that 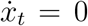 when *x* = *σ v/κ* for the harmonic potential *U*(*x*) = *κx*^2^/2. Without confinement (*U* = 0) and with little thermal noise (*β* = 10), the trajectories look as plotted in Fig. 2(a). That is giving rise, as for *β*_eff_ = 0, to edge states; compare Fig. 2(a) with Fig. 2(b) which represents trajectories in the standard RTP-process for which *T* = 0, *β*_eff_ = 0.

**FIG. 2:**
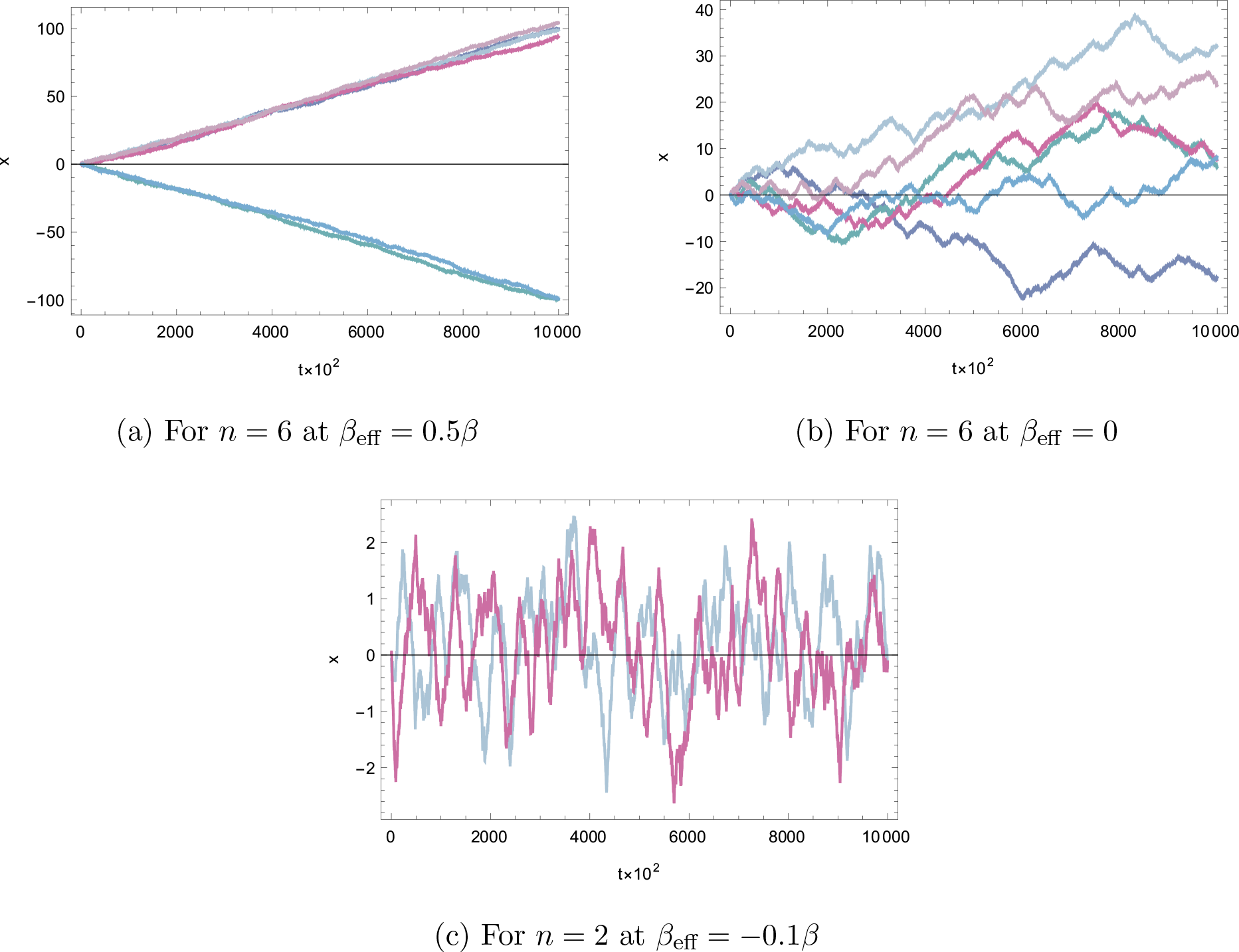
Trajectories of *n* particles under free low-temperature locomotion (II.1)–(II.2) with *U* = 0, *v* = 1, *β* = 10, *α* = 0.5, for different *β*_eff_, from low(a) to high(b)–(c) internal temperatures.

Suppose next that *β*_eff_ *<* 0. The hot spot has reached a negative temperature (indicating population inversion for a multilevel system). The flipping rate *α* exp{−*β*_eff_ *v σ x}* is going to be small when *σ x <* 0, and large when *σ x >* 0. Hence, if initially the particle is pushed with *σ* = 1 to the positive *x >* 0, it will flip a lot and its motion remains bounded. Similarly, when *σ* = −1 and pushed to negative *x <* 0, it will flip a lot, and return to the origin. That implies that for *β*_eff_ *<* 0 the particles remain confined around the origin even for relatively small *α*, and even when *U* = 0 (no confining potential) at small *T*. The typical trajectories (for *U* = 0, *β* = 10, *β*_eff_ *<* 0) are shown in Fig. 2(c). For *β*_eff_ ≃ *β*, we need a large flipping rate to see the (passive) confined regime, as illustrated by comparing Fig. 3(a) and (b) with Fig. 3(c). See Fig. 4 for the boundary of the two regimes.

**FIG. 3:**
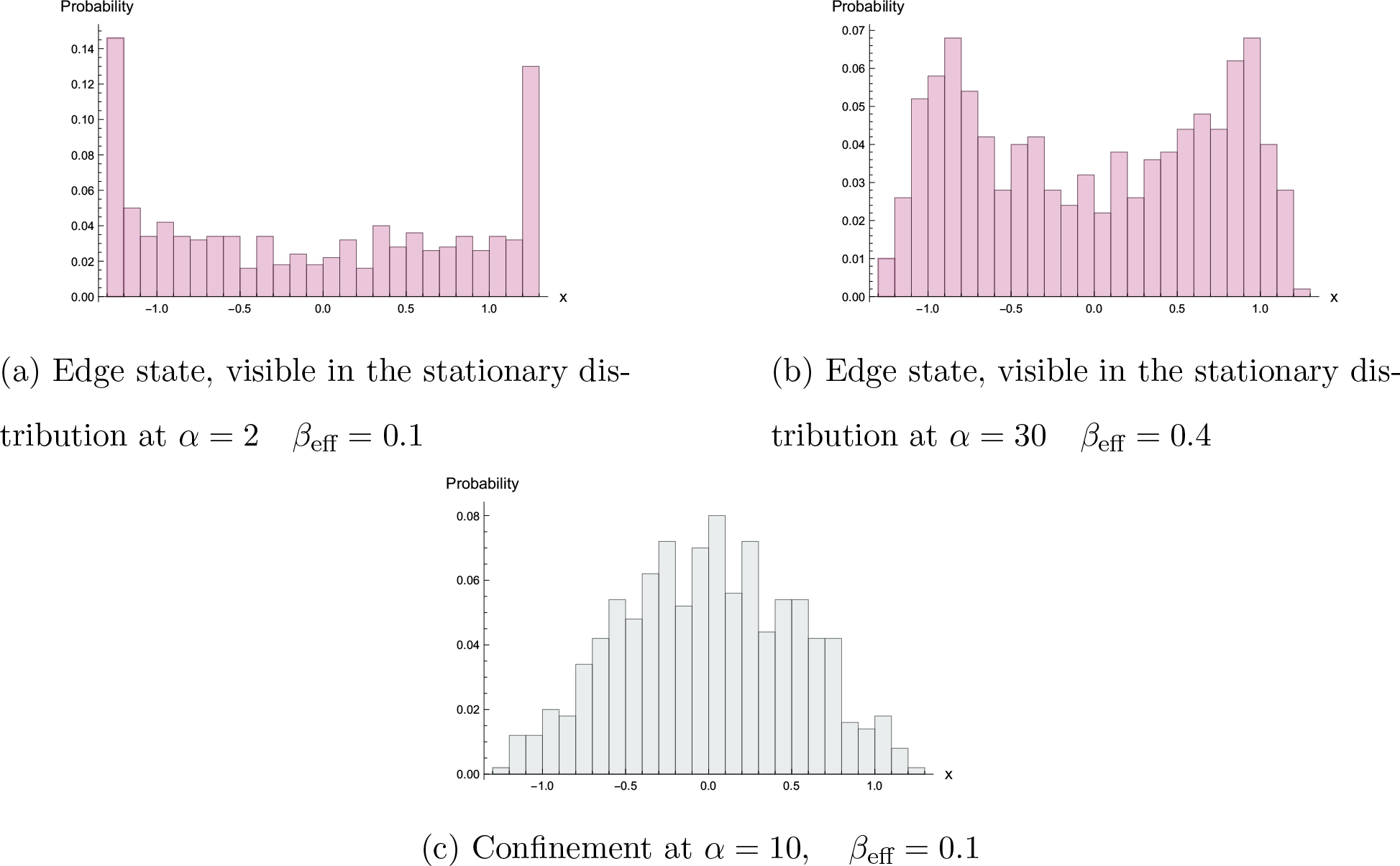
Stationary density of the two-temperature RTP-process at *v* = 3, *T* = 0 with confining potential *U* = *x*^2^. For large enough *α*, depending on *β*_eff_ and *v*, the two peaks at the edges merge and we enter the confined regime with a single peak at the origin.

**FIG. 4:**
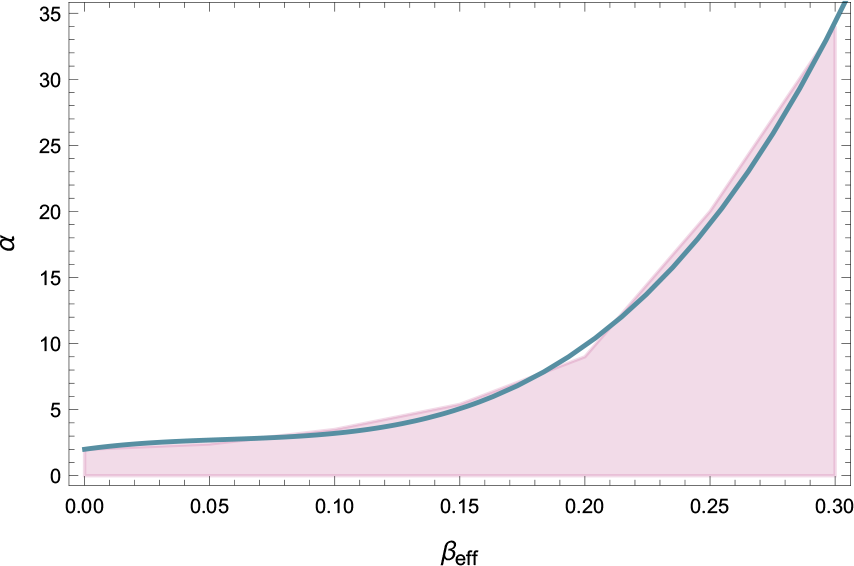
Shape transition in the (*β*_eff_, *α*)-plane for harmonic potential *U* (*x*) = *x*^2^ at *T* = 0 and with propulsion speed *v* = 3. The colored section represents the edgy regime. For high enough *α* starts the confined regime, with a sharply increasing transition line, fitting the transition as observed numerically by coloring 40 points depending on the concavity (colored) *versus* convexity of the simulated stationary density at the origin; see also Fig. 3.

For *β*_eff_ *> β*, we would have that the hidden bath has a lower temperature than the environment, which would make changes in configuration less efficient (as if the biological process must warm up the hidden degrees of freedom, instead of using their energy).

#### 2. Heat

The parametrization in the two-temperature process (II.1)–(II.2) easily identifies the driving and permits a standard discussion of thermal properties. Since the system is open (to two baths) the energy *E*(*x, σ*) = −*vσx* + *U*(*x*) changes and it does so in two ways:

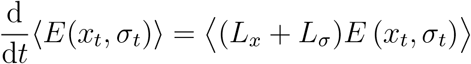

for

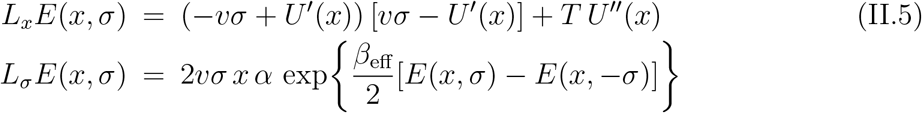

Therefore, upon specifying the instantaneous variables (*x, σ*),

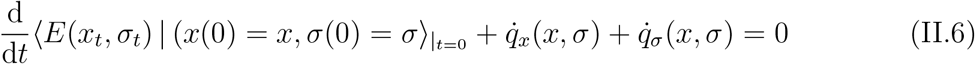

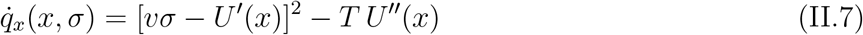

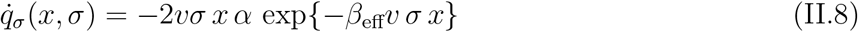

where 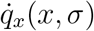 equals the expected heat flux to the ambient bath at inverse temperature *β*, and 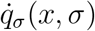 is the expected heat flux to the internal bath at inverse temperature *β*_eff_. Of course, in the steady distribution, their sum 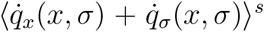 (expected value in the stationary distribution) equals zero, and there is a constant flow of energy between the two heat baths. From the above, as is common practise now by using standard methods such as outlined in [27, 31, 32], various thermal properties can be derived such as variable and mean entropy production, heat capacities, and delivered work in heat conduction networks. We only sketch, next, the less common example of constructing the McLennan ensemble, [33, 34].

#### 3. McLennan regime

Thanks to the two-temperature embedding, we are able to spell out the *feverish* close-to-equilibrium regime, where *T* gets closer to the effective temperature *T*_eff_.

We come back to the case (II.1) with confining potential and thermal noise. We are interested in the stationary distribution for its first (linear, nonequilibrium) correction to (II.3).

Imagining that (*β* − *β*_eff_) *v x* remains small (no big excursions), the considered backward generator for (II.1) has the form *L* = *L*_eq_ + *δ L*_1_ where the perturbation *δ* = 1 − *β*_eff_*/β* is in the flipping part, giving the dissipated power (A.1),

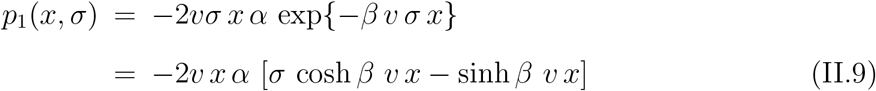

which can be inserted in the general formulæ (A.2)–(A.3), and from which we obtain the McLennan distribution *ρ*_ML_ = *ρ*^eq^(1 − *βW*), with *W* (of order *δ*) from (A.2).

We already see however from (II.9), and as a good approximation to the effective potential for small *α*, that for *U*(*x*) = *x*^2^/2, the potential 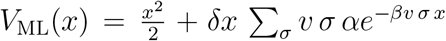 may generate an edge state: we refer to Fig. 5, where *V*_ML_(*x*) is plotted, with a minimum in *x* = 0, and going to −∞ for *x* → ∞. The calculated dependence on *α* is as already seen in Fig. 4; there are edge states arbitrarily close to equilibrium at very low temperatures.

**FIG. 5:**
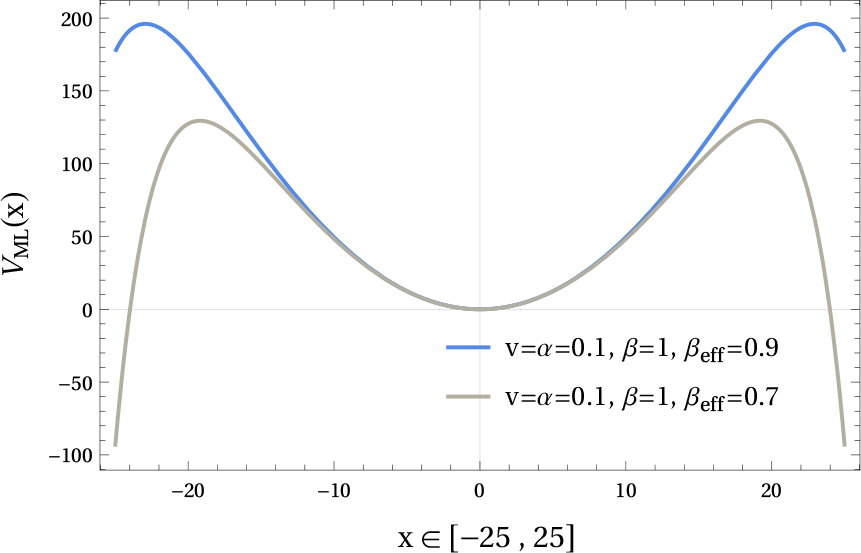
The effective potential generating the close-to-equilibrium ensemble, showing the occurrence of edge states as already observed in Figs. 3–4.

### B. Active Brownian motion

We move to two dimensions with position 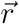 for a point particle, satisfying the equation of motion in Cartesian coordinates 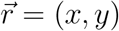,

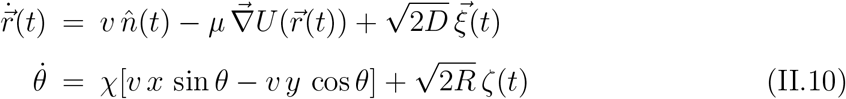

where the unit vector 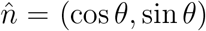 replaces the dichotomous noise in (II.1). As before, 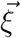 is standard white noise on the two-dimensional locomotion of the particle, while *ζ*(*t*) is rotational white noise, constantly redirecting the particle. The external potential *U* is confining and *μ >* 0 is a mobility coefficient. The constant *χ* can be taken positive or negative. Note that we still deal with a point particle and the *θ*−dynamics is indicating the direction of locomotion.

We basically repeat the same remarks and analysis of the previous section on RTPs. The (*x, y*)−dependence in (II.10) is not visible in the locomotion (first equation for 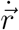) and refers to a spatial frame where the origin is determined by the confining potential *U*. The ambient temperature *T* and the internal effective temperature *T*_eff_ follow from

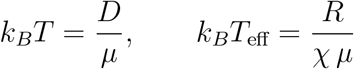

For *χ* = 0 we get the standard active Brownian process, which, in the present setup, refers to infinite effective temperature *T*_eff_ = ∞ (as it was for standard run-and-tumble particles as well); see *e.g*., [35].

In our setup and with a confining potential *U*, we imagine an energy function

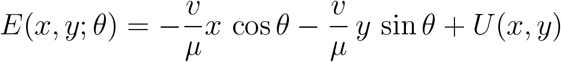

coupling the hot spot (with coordinate *θ*) with the spatial coordinates (*x, y*). There is *a priori* no problem with taking *T*_eff_ *<* 0 via *χ <* 0 as that temperature is effective and may refer to population inversion.

There are again two equilibrium cases. The persistence length is ℓ = *v/R*, reducing the dynamics to passive (or usual) Brownian motion in the limit ℓ ↓ 0. Secondly, when *Dχ* = *R* we get *T* = *T*_eff_ and detailed balance gets restored, with equilibrium density (for *U* confining)

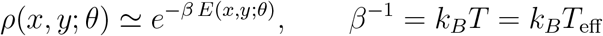

We can proceed from this case, by varying *χ*, and basically repeating the same analysis as in Section II A for run-and-tumble particles. The first dynamical phase is when *χ* ≤ 0, where the system is unconfined when *U* = 0 see Fig. 6, and the second phase happens for *χ >* 0 where the dynamics is confined, even when *U* = 0, see Fig. 7. Furthermore, the notion of heat and the characterization of the close-to-equilibrium regime is clear, as in the previous sections for the RTPs, since local detailed balance is verified for all *χ*.

**FIG. 6:**
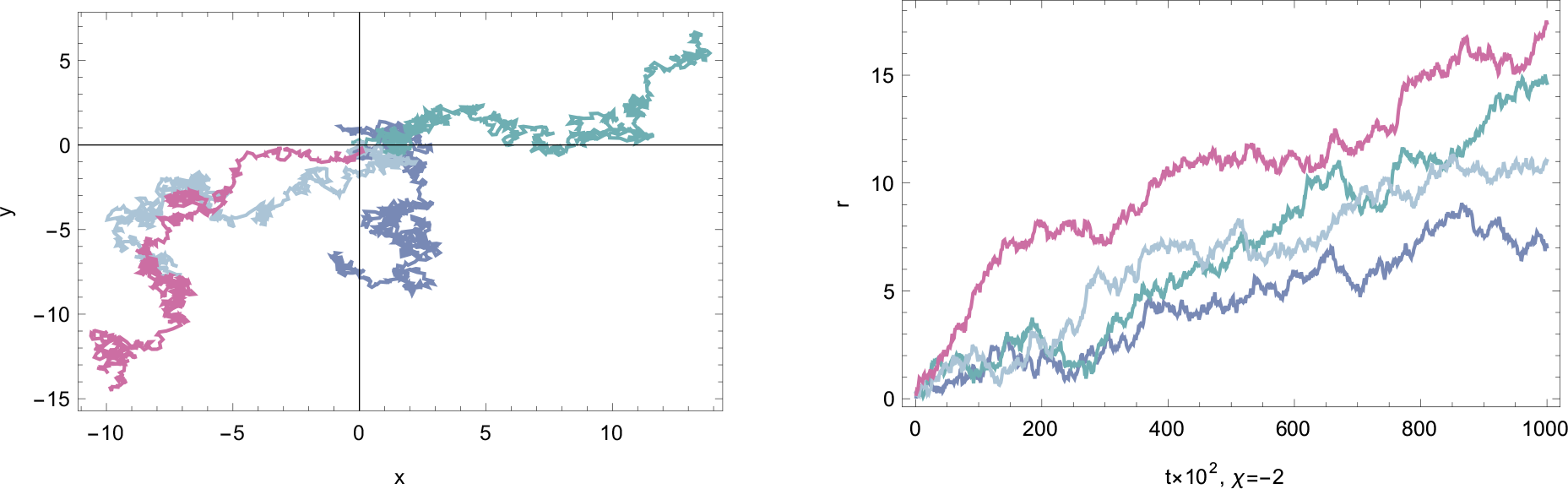
Active Brownian motion for four particles with the dynamics (II.10) at *χ <* 0, when *v* = *R* = *D* = 1 and *U* = 0. Left: trajectories when the particles start from *r*(0) = 0. Right: the distance of the particle positions from the origin.

**FIG. 7:**
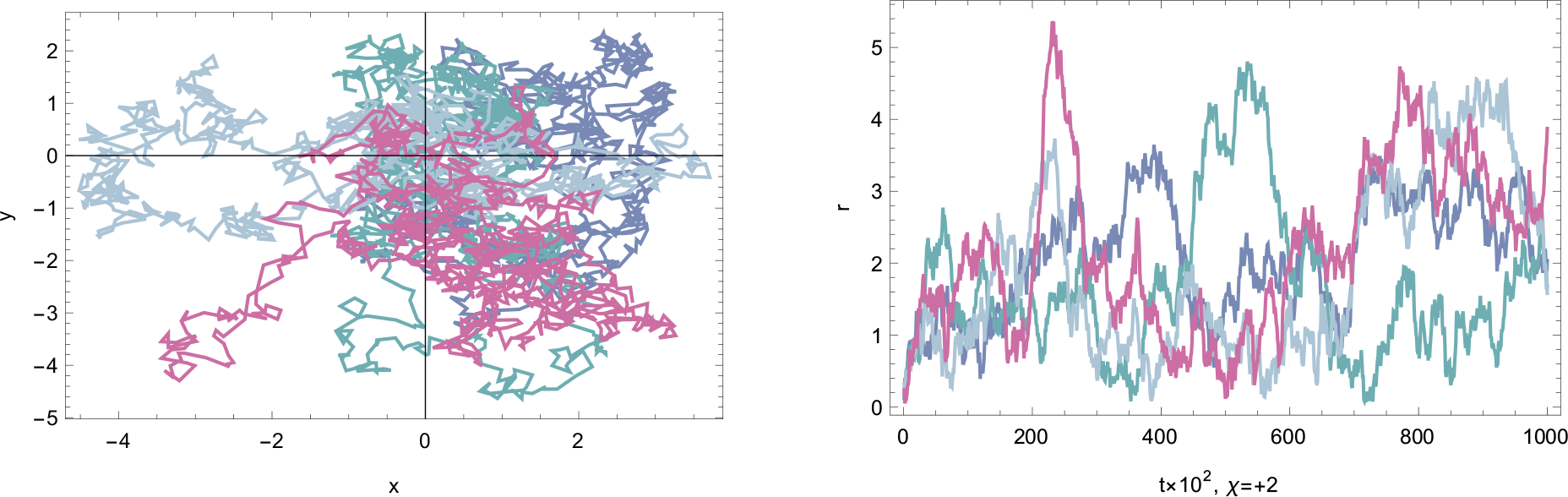
Active Brownian motion for four particles for the dynamics (II.10) with *χ >* 0, when *v* = *R* = *D* = 1 and *U* = 0. Left: trajectories when the particles start from *r*(0) = 0. Right: the distance of the particle positions to the origin.

## III. HOT SPOTS AND EFFECTIVE TEMPERATURE

The issue of temperature and energy fluctuations in mesoscopic systems subject to nonequilibrium processes is a topic of much interest; see *e.g*. older and recent references in [36]. To associate an effective temperature, and especially a high temperature, to specific agents or areas within a living system obviously raises even more questions. It is to be understood in what sense that temperature is measurable and compatible with other constraints, *e.g*., avoiding high–temperature gradients and damage caused by a local high temperature. Whether it can be measured (*e.g*. by fluorescence techniques) is not clear, and in what sense it is at all compatible with the stability of the morphology is also an open question. However, since some good time, the presence of “hot spots” has been widely discussed in the specialized literature [12–17], while the jury is still out for deciding on the precise nature of what is experimentally seen. The subject and scope of the present paper are clearly much more theoretical, *e.g*. for allowing a consistent discussion of heat in active particle models, using the coupling with a hot spot. Here, we add two examples of such spots, the first being a Markov jump model of an agitated molecule (making it a molecular switch), and the second a randomly agitated vibrational mode, where we specify the effective temperature as a measure of the width of the population homogenization with respect to the equilibrium Boltzmann-Gibbs statistics.

### A. Multilevel switch

As an example of an agitated molecule, we consider a switch. Molecular switches [37–39] are biologically involved in signaling pathways. They are proteins that flip between two architectures causing either an *on*- or *off*-state for intracellular signaling. The *on* is an active state, typically initiated from an external stimulus, [40]. Other versions include genetic toggle switches, which in turn may have practical applications as a synthetic addressable cellular memory, [41].

One type of modeling proceeds via specifying a multilevel system where, *e.g*. at random times, energy levels are shifted. The change in energies may be provoked by the (sudden) entrance of an ion, a proton, or even a photon that modifies the molecular configuration or the harvesting of energy on even smaller (nano)scales. Such processes happen on the level of individual molecules.

A simple (and more extreme) model of a molecular switch has its energy ladder inverted at random moments in time. We imagine an energy labeled by a coordinate *η*, which is mirrored by flipping a variable *σ* indicating the *on*- or *off*-state; see Fig. 8(a). More precisely, we consider energies

**FIG. 8:**
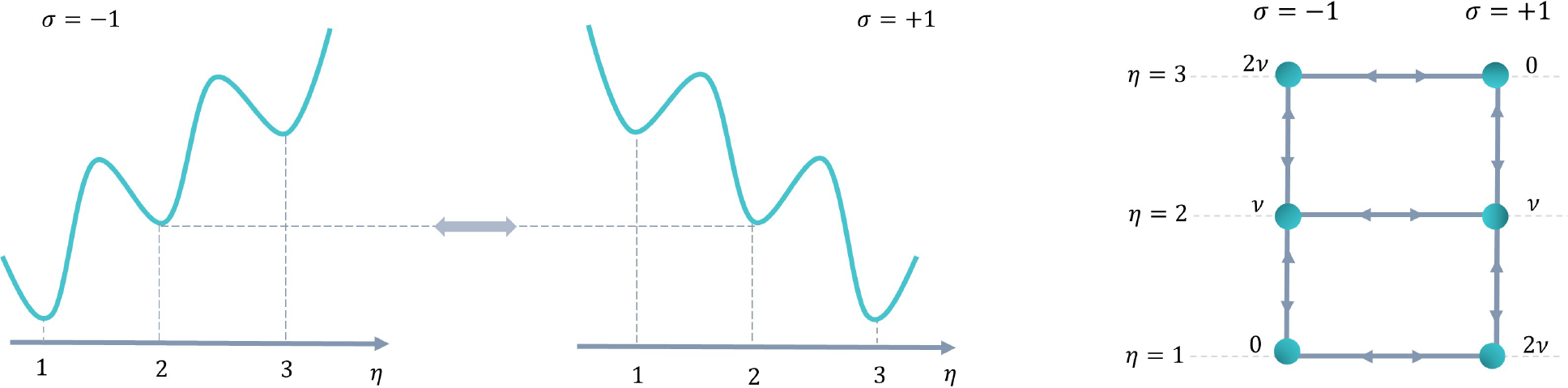
Modeling a three-level switch (*n* = 3). Left: switching the energy landscape. Right: graph of transitions with an indication of energy.

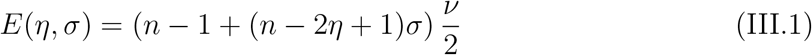

separated by energy unit *ν >* 0. For each *σ* = ±1, there are energy levels *η* = 1, …, *n* for a fixed integer *n* ≥ 2. Typically, *n* can be considered small. The symmetry *E*(*η*, −*σ*) = *E*(*n* − *η* + 1, *σ*) implies that by flipping *σ* the energy ladder is simply reversed. That switch will be responsible for a nonequilibrium occupation statistics. For *n* = 2 we are dealing with a switch for a two-level system (or classical qubit) where at random times the ground state and the excited state get exchanged, [21, 32].

Transitions between energy levels are described by a Master equation, for which we introduce transition rates *k*_*σ*_(*η, η* ± 1) (for moving up or down in energy at given *σ*), and there is a constant rate *k*_*η*_(*σ*) = *α* for switching *σ* → −*σ* at a given *η*. We refer to Fig.8. (b) for a graphical representation for *n* = 3. Of course, at *η* = 1 and *η* = *n*, the system can only move up or down the ladder (or switch).

In particular, we take the parameterization,

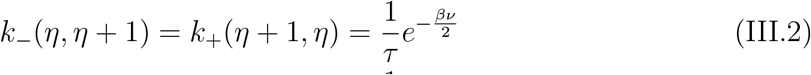

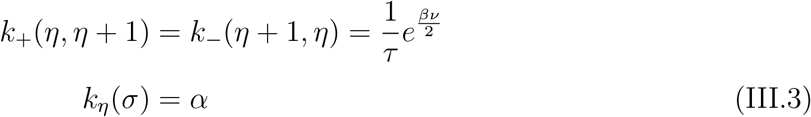

where *ατ >* 0 indicates the switching amplitude compared to moving up/down the ladder in Fig.8). Units of time are arbitrary here, as we only deal with the steady condition. We have also not specified the kinetic barrier in terms of the time-scale *τ* in the energy transitions (III.2); that would introduce another dependence on *β* which we skip for the present discussion. Finally, we imagine many independent copies of that system, which makes the analysis meaningful.

In the limit *α* ↓ 0, one disconnects the two sides of the ladder obtaining two separate equilibrium systems. They are described by the Boltzmann–Gibbs distribution *ρ*_eq+_(*η*) ∼ exp −*β*(*n* − *η*)*ν* and *ρ*_eq-_(*η*) ∼ exp −*β* (*η* − 1)*ν* which are of course identical by a simple change of variable. Without agitation, the thermal properties of the molecule are just that of an equilibrium system at inverse temperature *β*.

In order to estimate the effective temperature of the system we keep the energy-variable *η* coupled to the ambient bath but we also parameterize the switching in (III.3) via

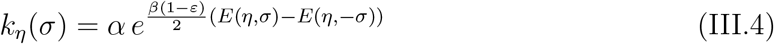

for some *ε* ≥ 0. Detailed balance holds true for *ε* = 0. A the other end is the usual setup for switching, where one would take *ε* = 1 and thus have a (purely) random flipping between two energy ladders (as in (III.3)). Then, the switch occurs at rate *α* independently of the molecular state *η*, and no feedback occurs.

For *ε* = 1, as *α* ↑ ∞, we reach the uniform stationary distribution

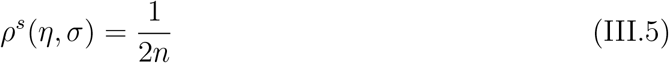

which can be considered the infinite-temperature distribution indeed corresponding with *β*′ = 0. See Appendix B for the proof.

Under (III.4). the *α* ↑ ∞-limit of the stationary distribution still depends on *ε*. As we move from *ε* = 0 to *ε* = 1, always for large *α*, the stationary distribution *ρ*^*s*^(*η*) over *η* becomes more and more uniform. In other words, the nonequilibrium driving adds weight to the levels that are less occupied in equilibrium, realizing effectively an occupation statistics corresponding to a higher temperature (infinite upon reaching (III.5) for *ε* = 1).

To be more specific we take the case of *n* = 2, and we define the effective temperature,

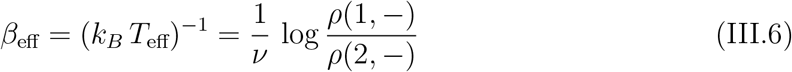

measuring the homogenization of energy occupation. In that case (of *n* = 2), there are four states and the stationary distribution is

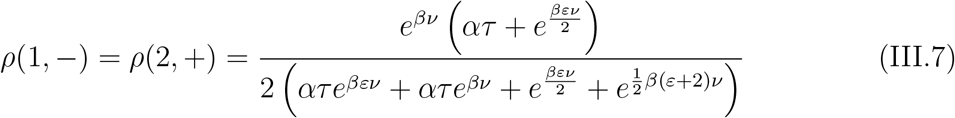

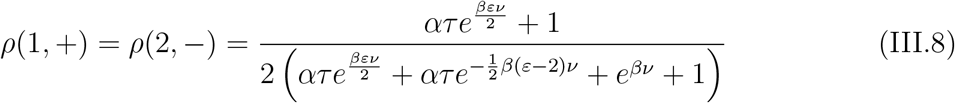

Those stationary probabilities are plotted in Fig. 9 for varying *ε* and *α*.

**FIG. 9:**
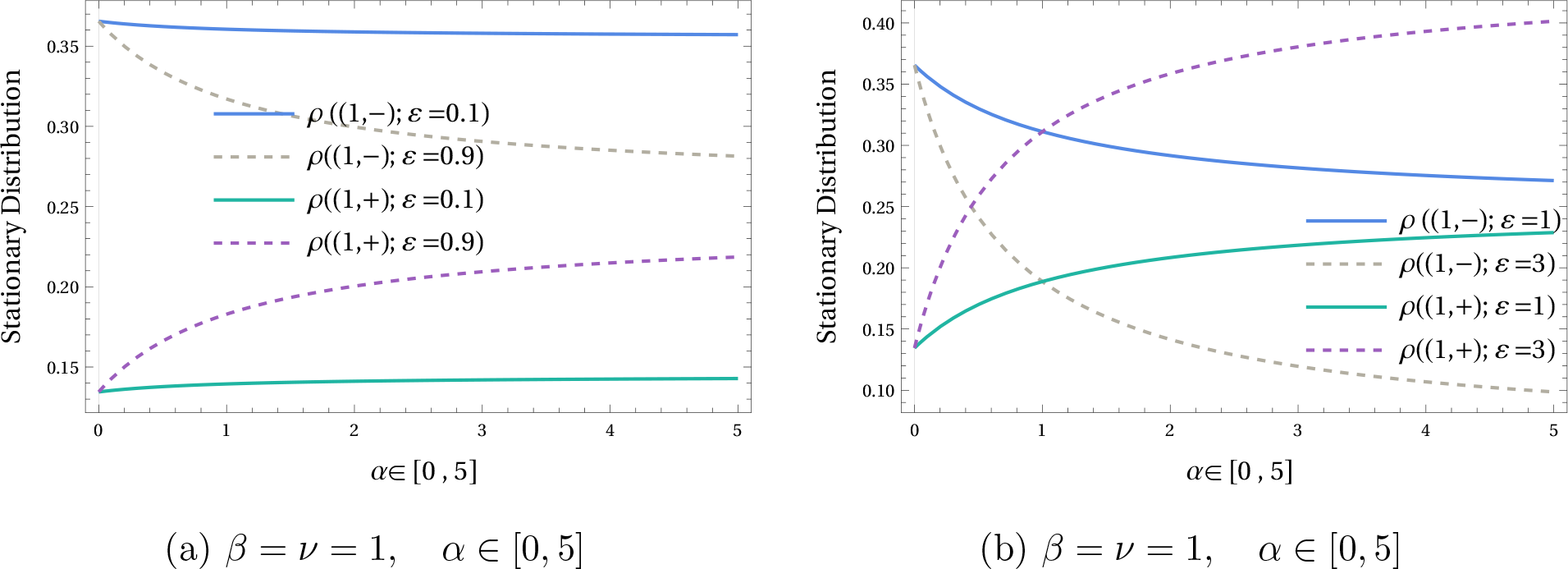
Comparing the stationary probabilities of states (1, +) = (2, −) and (1, −) for the dynamics with rates (III.2)-(III.4) for *n* = 2, (a) increasing *ε <* 1 brings the level probabilities closer, (b) increasing *ε >* 1 leads to population inversion, *e.g*. starting from *α* ≃ 0.2 for *ε* = 3.

Note that increasing *ε* uniformizes the stationary distribution for *ε <* 1, and then, for *ε >* 1 and at least for large enough *α*, leads to population inversion. It implies that the effective temperature (III.6) moves up from *T*_eff_ = *T* at *ε* = 0, to *T*_eff_ = ∞ at *ε* = 1, and then becomes negative for *ε >* 1 (for large enough switching rate *α*); see Fig. 10 qualifying the model as describing a hot spot for *ε* ≃ 1.

**FIG. 10:**
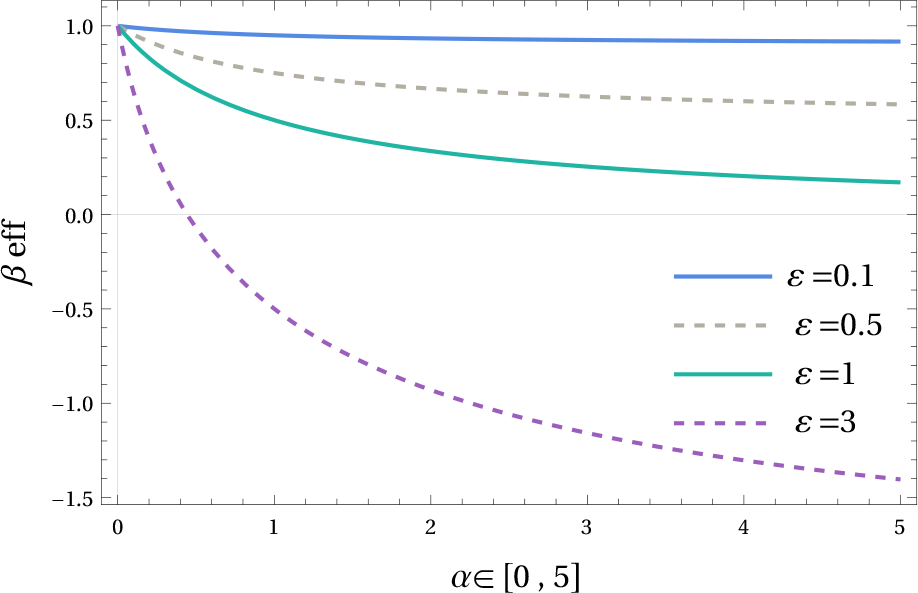
The effective inverse temperature (III.6) for *β* = *ν* = 1, as function of *α* ∈ [0, 5].

### B. Agitated vibrational mode

A toy model that has been used for generating motion of particles is that of Brownian ratchets, [42–50]. The swimming is made possible by exploiting or creating a time-dependent landscape. We represent that landscape again by a variable *η*, as denoting the energy levels in the molecular switch of the previous section, but now it takes continuous values and may indicate the value of a vibrational mode in a molecule.

A simple example already appeared in [20, 51] for illustrating active calorimetry, and is obtained by taking a harmonic potential Φ on the real line with a spring ‘constant’ *κ*(1 ±*w*) switching values,

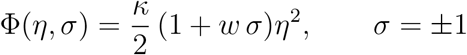

We take *κ >* 0, but *w* can be negative and even have |*w*| *>* 1 to allow higher activation. The idea is that *κ η*^2^/2 is still measuring the vibrational energy. The binary variable *σ* flips at rate *α >* 0, while

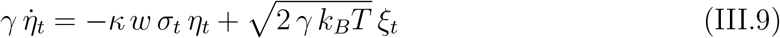

As for an overdamped dynamics in a background at temperature *T*, we take *ξ*_*t*_ standard white noise, and we put *γ* = 1, *k*_*B*_ = 1 as before.

To get a measure of the effective temperature, we take the spreading ⟨*η*^2^⟩^*s*^ in the stationary distribution (if it exists). It indeed senses the occupation at large energies. In equilibrium (*w* = 0), we have ⟨*η*^2^⟩^eq^ = *T/κ*.

It is easy enough to obtain the moment equations from (III.9),

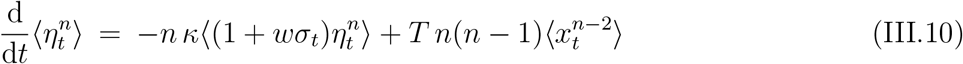

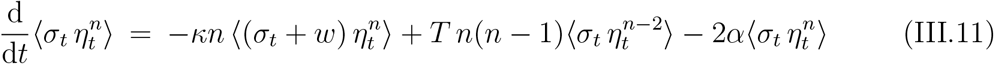

The stationary distribution is nonGaussian in general. Yet, putting the above equations equal to zero, we find that the stationary moments *a*_*n*_ = ⟨*η*^*n*^⟩^*s*^ and *b*_*n*_ = ⟨*ση*^*n*^⟩^*s*^ verify recurrence formulæ

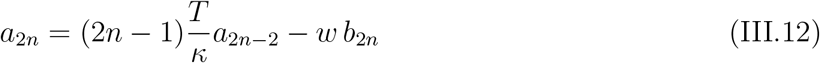

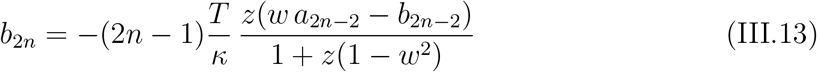

in terms of the dimensionless persistence *z* = *κ/α*.

When |*w*| *>* 1, there is a critical persistence

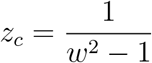

and for *z > z*_*c*_ there is no stationary distribution, [51]. In particular, the second moments equal

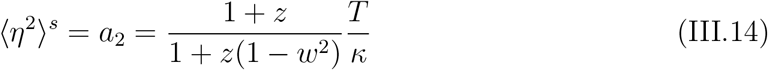

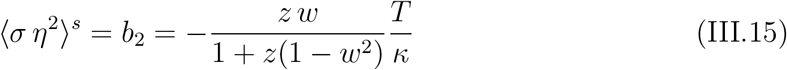

Of course, we see the usual equilibrium for *w* = 0, with a deviation, for *α >* 0, governed by 1 − *w*^2^. When |*w*| *<* 1, the effective temperature is slightly larger than *T*. When |*w*| *>* 1, the stationary second moment ⟨*η*^2^⟩^*s*^ diverges for *z* ↑ *z*_*c*_, which means that the effective temperature reaches infinite as the flip rate *α* ↓ *α*_*c*_ = *κ* (*w*^2^ − 1). Then indeed, the system increasingly occupies very high energy states in that single vibrational mode. It has become a “hot spot” indeed.

## IV. CONCLUSIONS

Life on our planet has been running on the temperature difference between the Sun (hot spot of low entropy) and vast outer space. ATP and other chemical fuels appear to recreate that nonequilibrium force inside organisms on Earth, activating many (but tiny) open systems. They violate detailed balance, and effectively mimic a very-high-temperature occupation statistics. When coupled to other degrees of motion at a different ambient temperature, a two-temperature heat engine is created (though needing to work much faster than allowed for in the Carnot regime).

Apart from that conceptual framework, which is generally useful for understanding efficiencies in active or bio-mimicking matter in material science [2], the embedding of active particle models in two-temperature processes allows the use of statistical thermodynamics applied to small systems. Local detailed balance gets restored, and a clearer prescription of heat and entropy production becomes possible, witnessed *e.g*. by including a description of the close-to-equilibrium (McLennan) regime for active particles.

The present paper has illustrated these points using simple models of bacterial locomotion (run-and-tumble and active Brownian particles), and by providing examples of hot spots (molecular switches and agitated vibrational modes). The two-temperature embedding of the run-and-tumble process has in addition shown the robustness of the shape transition between a more confined regime and the occurrence of edge states, as monitored by the tumbling rate.

## Acknowledgments

FK thanks Simon Krekels and Kasper Meerts for fruitful discussions on the subject of RTPs and for help with the numerical work. CM thanks Gianmaria Falasco for pointing to relevant references.

## Appendix A McLennan distribution

To show how to be quantitative about the distance to equilibrium, we discuss the regime close-to-equilibrium and how it retains a more universal interpretation. Indeed, identifying the so-called McLennan ensemble governing the close-to-equilibrium regime is not just a perturbative result. Obviously, it is part of linear response, but it gets more interesting when there is a physical meaning associated to that ensemble. In the same sense that the Gibbs formalism comes with extra interpretation and can start from knowing some basic physical quantities (where the Hamiltonian is of central importance), the McLennan ensemble can be constructed “physically”: its central quantity is the transient entropy flux, but it does require understanding the thermal structure of the dynamics. To obtain the McLennan ensemble for active processes is an application of the program described in Section II.

We remind the reader about the close-to-equilibrium ensemble when such a regime is possible to be defined. It starts from a clear notion of an equilibrium reference dynamics in weak contact with a thermal environment at uniform inverse temperature *β*, for which equilibrium can be defined and studied via the Boltzmann-Gibbs distribution *ρ*^eq^(*x*) ∼ exp −*βE*(*x*) for energy function *E*. Next, a driving is applied in the spirit of local detailed balance; see *e.g*. [26]. That leads to identifying the transient irreversible entropy flux. More specifically, we have a driving amplitude *ε* and we find the expected dissipated power 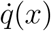 when in state *x*. We need its linear order in the perturbation, of strength *δ*

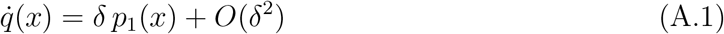

to build the work function

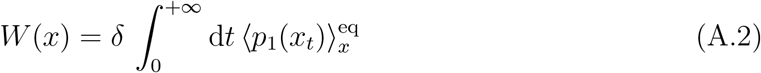

where the expectation 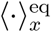 is in the equilibrium dynamics for dynamical variable *x*_*t*_ (which is left unspecified here), starting from *x*_0_ = *x*. The mean entropy change per *k*_*B*_ in the heat bath is given by *βW* (*x*) when starting the equilibrium dynamics from the system in state *x*. It is exactly that (finite) transient component of the irreversible part of the entropy flux that corrects the Boltzmann distribution to first order in the driving: we define the McLennan quasipotential

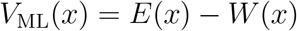

where *E* is the energy, and the McLennan distribution is

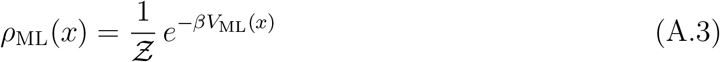

For its thermal properties, the heat capacity in the McLennan ensemble was recently studied in [52]. That distribution is called after McLennan because of the introduction in [34]; see also [33].

## Appendix B Proof of (III.5)

We prove the uniformization (III.5) by using a graphical representation of the stationary probability law *ρ*^*s*^. In general, the stationary solution of the Master Equation for a Markov jump process can be represented by the Kirchhoff formula

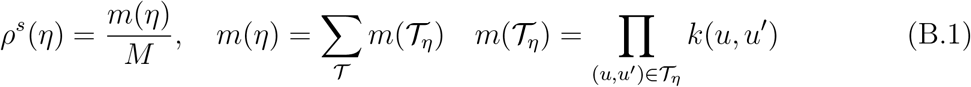

where *M* = ∑_*y*_ *m*(*y*), 𝒯 is the set of all spanning trees, *𝒯*_*η*_ is a spanning tree rooted at *η* (all edges are directed toward *η*) and *m*(𝒯_*η*_) is the weight of the directed spanning tree 𝒯_*η*_; see [53, 54].

Consider a ladder with *n* steps and with rates given in (III.2)–(III.3), where *ε* = 1. In the limit *α* ↑ ∞, (B.1) is dominated by the contribution of spanning trees passing over all “switching edges.” For example, for the ladder with *n* = 3 steps, the dominating spanning trees are shown in Fig. 11.

**FIG. 11:**
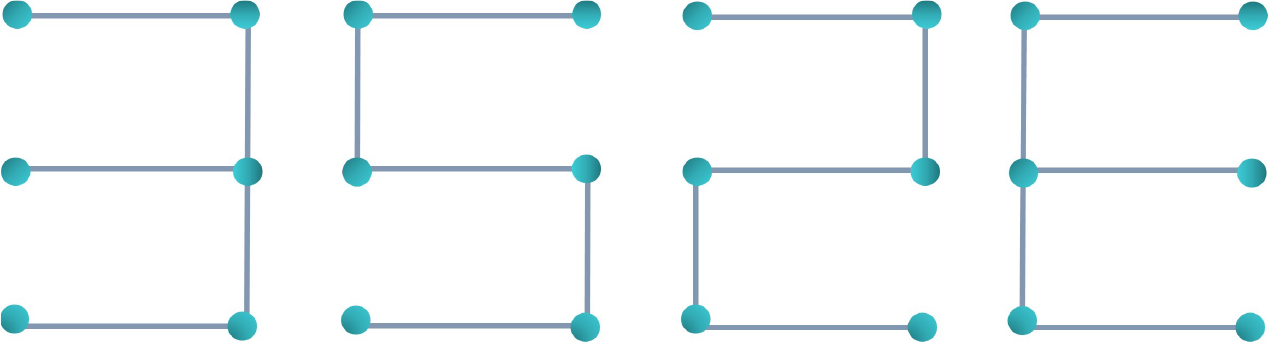
Nonoriented spanning trees of the ladder *n* = 3, that dominate in the Kirchhoff formula.

We start by showing the symmetry, when *α* ↑ ∞, for all *η*,

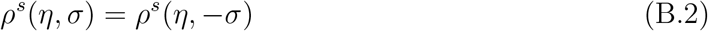

For every tree rooted in (*η, σ*) there exists a tree with the same weight rooted in (*η*, −*σ*). The new tree is created by changing the direction of the edge (*η*, −*σ*) → (*η, σ*) into (*η, σ*) → (*η*, −*σ*); see Fig. 12.

**FIG. 12:**
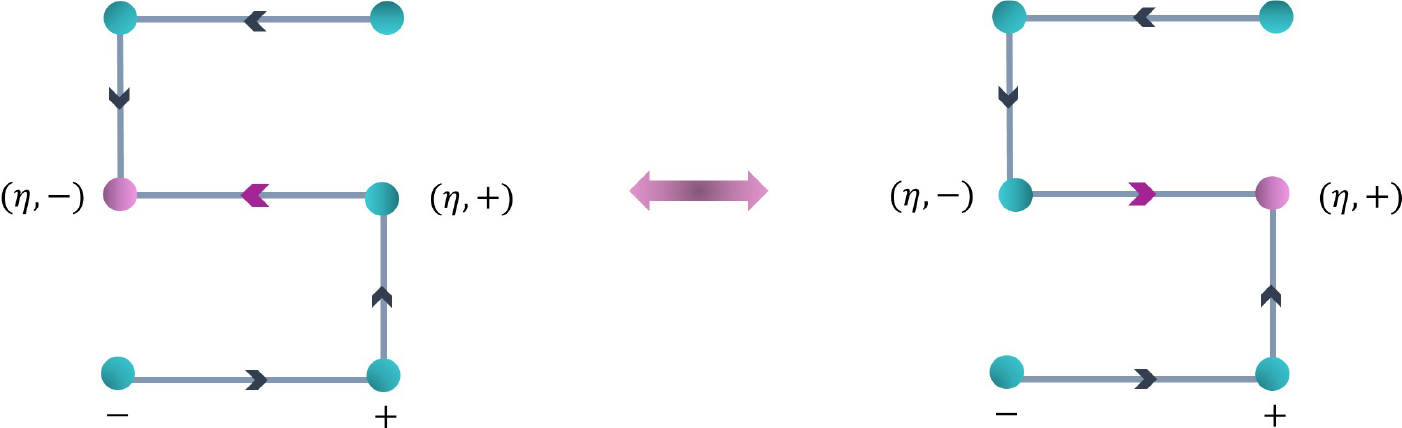
For every rooted tree, a new tree can be created by changing the direction of one edge. This new tree will be rooted in the target of the edge’s new direction. By changing the direction of the red edge, a new spanning tree rooted at (*η*, −*σ*) is created.

As a consequence, the sum over all weights of the dominating trees in *ρ*^*s*^(*η, σ*) and *ρ*^*s*^(*η*, −*σ*) are equal; see(B.1).

Next we show that when *α* ↑ ∞, for all (*η, σ*),

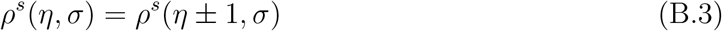

Take a tree rooted at (*η, σ*). There is a vertical edge which is connecting *η*±1 to *η*. That edge can be in *σ* or −*σ*. By replacing the edge to the opposite *σ*, and changing the direction, the weight does not change. By changing the direction of a horizontal edge, the weight remains the same as well. Hence, for every tree rooted at (*η, σ*), there exists a tree with the same weight rooted at (*η* + 1, *σ*) such that the direction of one horizontal edge and of one vertical edge are changed; see Fig. 13.

**FIG. 13:**
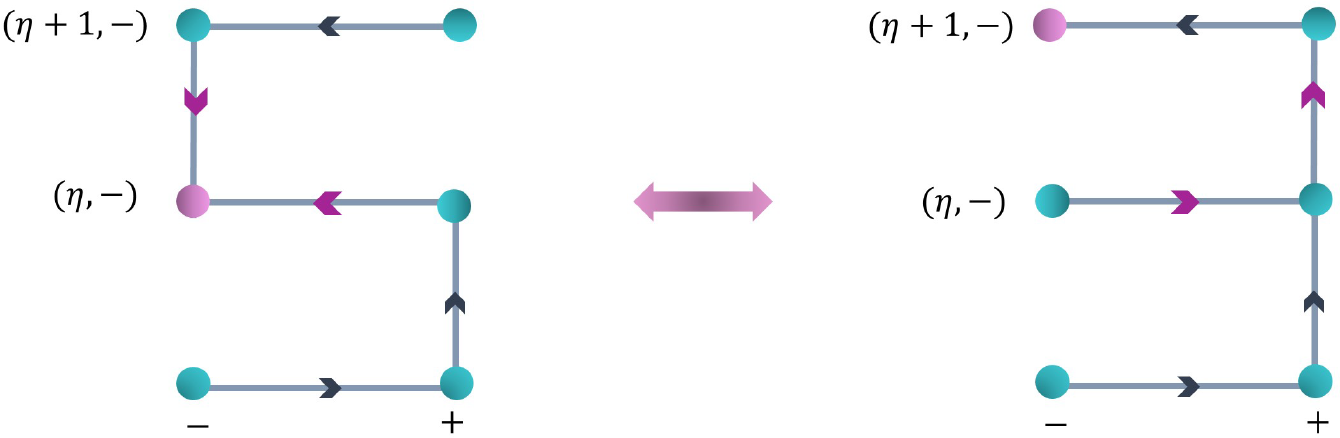
By changing the direction and *σ* of a vertical edge and changing the direction of a horizontal edge, the weight of the tree is not changing. Left: a spanning tree rooted at (*η, σ*), Right: a new spanning tree with the same weight rooted at (*η* + 1, *σ*).

Finally, from (B.2) and (B.3) we see that all states have the same stationary probability, 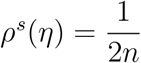.

## Notes

### Competing Interest Statement

The authors have declared no competing interest.

### Summary of Updates

caption of the figure 1 is changed

